# Clinical Phage Microbiology: A suggested *in-vitro* framework for phage therapy

**DOI:** 10.1101/2021.02.23.432393

**Authors:** Daniel Gelman, Ortal Yerushalmy, Shira Ben-Porat, Chani Rakov, Sivan Alkalay-Oren, Karen Adler, Leron Khalifa, Mohanad Abdalrhman, Shunit Coppenhagen-Glazer, Saima Aslam, Robert T Schooley, Ran Nir-Paz, Ronen Hazan

**Affiliations:** Institute of Dental Sciences, School of Dentistry, Hebrew University of Jerusalem, Jerusalem, Israel; Department of Military Medicine, Faculty of Medicine, The Hebrew University of Jerusalem, Jerusalem, Israel; Hadassah-Hebrew University Medical Center, Department of Clinical Microbiology and Infectious Diseases, Jerusalem, and the Faculty of Medicine, The Hebrew University of Jerusalem, Jerusalem, Israel; Division of Infectious Diseases and Global Health, University of California, San Diego, La Jolla, California, USA; Center for Innovative Phage Applications and Therapeutics, University of California, San Diego, La Jolla, California, USA; Department of Medicine, University of California, San Diego, La Jolla, California, USA

## Abstract

Personalized-phage-therapy is a promising solution for the emerging crisis of bacterial infections that fail to be eradicated by conventional antibiotics.

One of the most crucial elements of personalized-phage-therapy is the proper matching of phages and antibiotics to the target bacteria in a given clinical setting. However, to date, there is no consensus guideline for laboratory procedures that enable *in vitro* evaluation of phages intended for treatment.

In this work, we suggest a framework and strategies identify appropriate phages and combine them with antibiotics in clinical microbiology laboratories. This framework, which we term here “Clinical Phage Microbiology” is based on our experience and other previously reported cases of both, successful and failed phage treatments.

Additionally, we discuss troubleshooting methodologies for possible pitfalls and special cases that may need to be assessed before treatment including interactions with the host immune system, biofilms, and polymicrobial infections.

We believe that the “Clinical Phage Microbiology” pipeline presented here should serve as the basis for standardization of laboratory protocols to match phages for personalized therapy.

## Introduction

Antimicrobial selection for the treatment of bacterial infectious diseases can be dictated by the antimicrobial susceptibility of bacteria isolated from patients, assessed by manual^1^ or automated methods^2^. The methodology of antimicrobial susceptibility testing has significantly evolved over several decades and is currently directed by the European Committee on Antimicrobial Susceptibility Testing (EUCAST) and the US-based Clinical and Laboratory Standards Institute (CLSI)^3^. This methodology is defined for each antimicrobial agent, to assure reliability and reproducibility in clinical practice.

Phage therapy, the use of phages as antimicrobial agents, has become a promising solution for persistent and resistant bacterial infections ^4,5^. A growing number of publications have described the clinical use of phage therapy, typically under compassionate regulation for investigational medical products ^6–11^. However, to date, there is no consensus on standard methodologies to evaluate phages for treatment. Phages were occasionally applied empirically, as shelf products with wide coverage, given without specific testing ^12,13^. In other cases, where personally selected treatments were given, the phages were selected by several different approaches, varying in their range and appropriateness for treatment design^6–10^. Thus, the need for consistent laboratory evaluation, from phage efficacy determination in a given clinical setting, through the design of suitable antibiotic combinations, to real-time adjustments, is highly important.

In this work, we suggest a preliminary pipeline, termed “Clinical Phage Microbiology”, as the basis for standardization of the laboratory protocols enabling clinical utilization of phage therapy. This pipeline aims to provide comprehensive and rapid testing of several aspects regarding the clinical use of phage therapy, to predict the best clinical outcome.

### Clinical Phage Microbiology Pipeline

The general view of the concept “Clinical Phage Microbiology” is presented in Fig 1. A key component of this approach is the development of large local phage collections, termed here “phage banks”, containing characterized^14,15^ phage preparations ^16,17^ ready for immediate clinical use^6,18^. The phage banks should be constantly expanded by screening for novel phages from raw samples^14^, by phage training, the adaptation of phages to specific bacterial isolates^19,20^, and by phage engineering ^21^.

**Fig 1.**
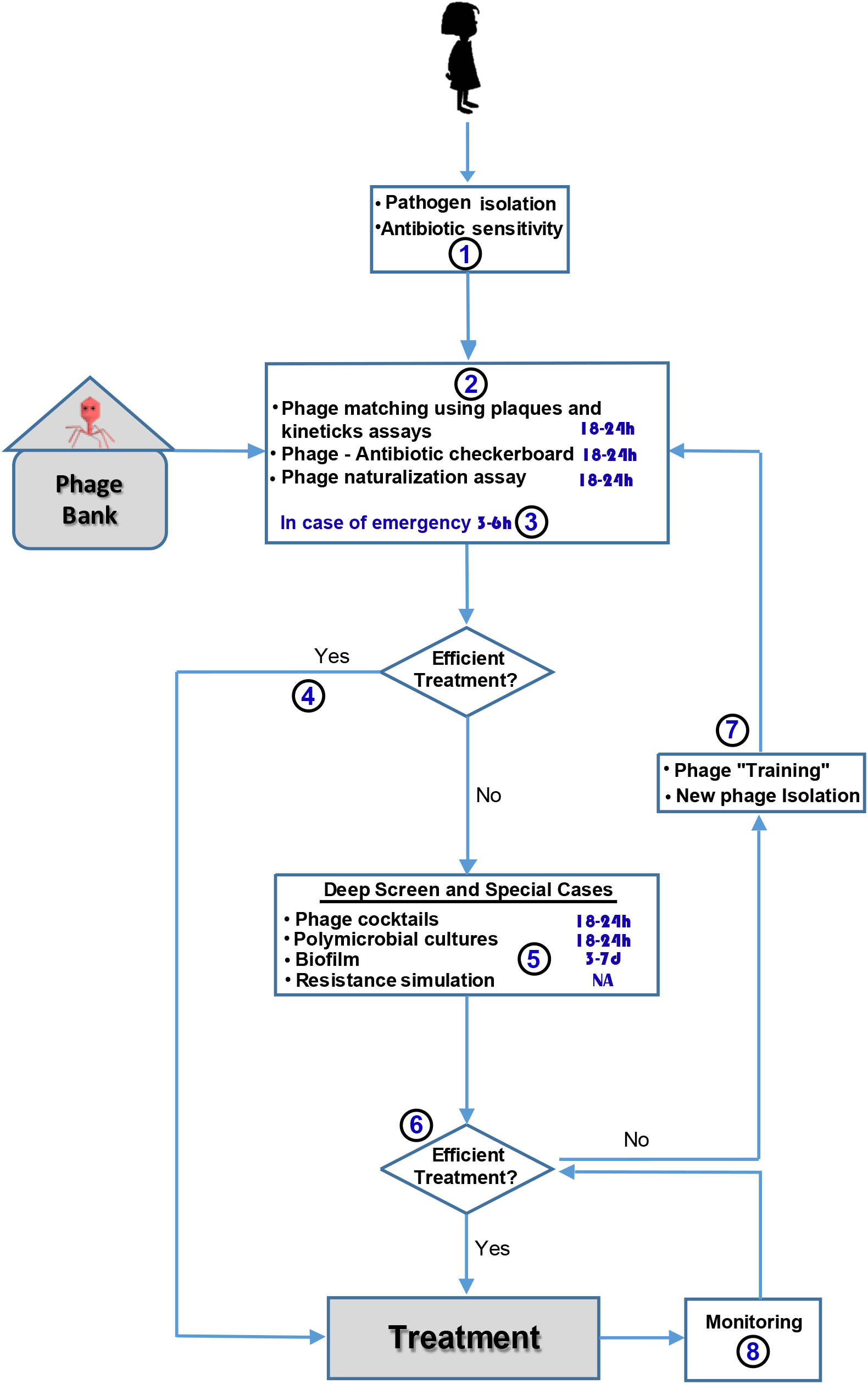
The Clinical Phage Microbiology Scheme. (1) Bacterial isolates from phage therapy candidates are identified and tested for antibiotic sensitivity. (2) The isolates are then screened against available phages from “Phage Banks” using several methods, including the plaque assay and growth kinetics monitoring. Appropriate phages are further assessed for suitable concomitant antibiotic regimens and potential host anti-phage immune response. (3) In cases of emergency, rapid matching may be performed instead. (4) If a suitable treatment regimen was found, the treatment may be initiated. (5) Otherwise, a preparation of phage cocktails with better coverage or a more comprehensive screen for suitable phage-antibiotic combinations should be pursued. Moreover, in clinical cases requiring special considerations, such as biofilms or polymicrobial infections, the intended treatment should be further evaluated inappropriate models, (6) and if found efficient, the treatment may be initiated. (7) If efficient treatment has not been found, additional phage isolation or phage training should be performed, followed by a new phage matching. (8) During treatment, constant monitoring for the emergence of resistant bacterial isolates and host interference to the treatment should be performed, followed by additional phage matching if required. Gray boxes are elements related to the scheme which are beyond the focus of the current work. In blue are estimated time frames for results obtaining with fast-growing pathogens.

The Clinical Phage Microbiology scheme begins with the isolation of bacterial pathogens from patients, followed by routine identification and antibiotic susceptibility testing^3,22^. Once the pathogen is isolated and identified, the next step of phage matching may be pursued.

### Matching Phages to Bacterial Targets

Accurate and rapid selection of the “appropriate” phages against target bacteria is an essential step of Clinical Phage Microbiology. Several previous published clinical trials have applied phages empirically-based only on theoretical activity^12,13^. However, these cases often led to treatment failure due to inappropriate phage selection. For example, in the case of the PhagoBurn trial, in which an empiric cocktail of twelve anti-*Pseudomonas aeruginosa* phages was used for the treatment of infected burn wounds, a clear association was found between the initial phage susceptibility of the clinical isolates to the rate of treatment success^12^. Thus, in our opinion, a targeted validation of phage efficacy is mandatory for precise phage therapy, and treatment should not be given without assessment of phage susceptibility.

#### A. Plaque Assay

One of the first and most established methods for phage efficacy determination^23^, which was also used in several clinical reports^8,21^, is the plaque assay (Fig 2A). In this method, phage suspensions are spotted onto bacterial lawns on agar plates, following the double-layer agar method^24^, and growth inhibition areas, or plaques, are evaluated^25^. Moreover, by spotting serial dilutions of a phage suspension, this method can be used to identify individual plaques and to determine the number of plaque-forming units (PFUs) of the phage on a specific bacterial strain^26^ (Fig 2B).

**Fig 2.**
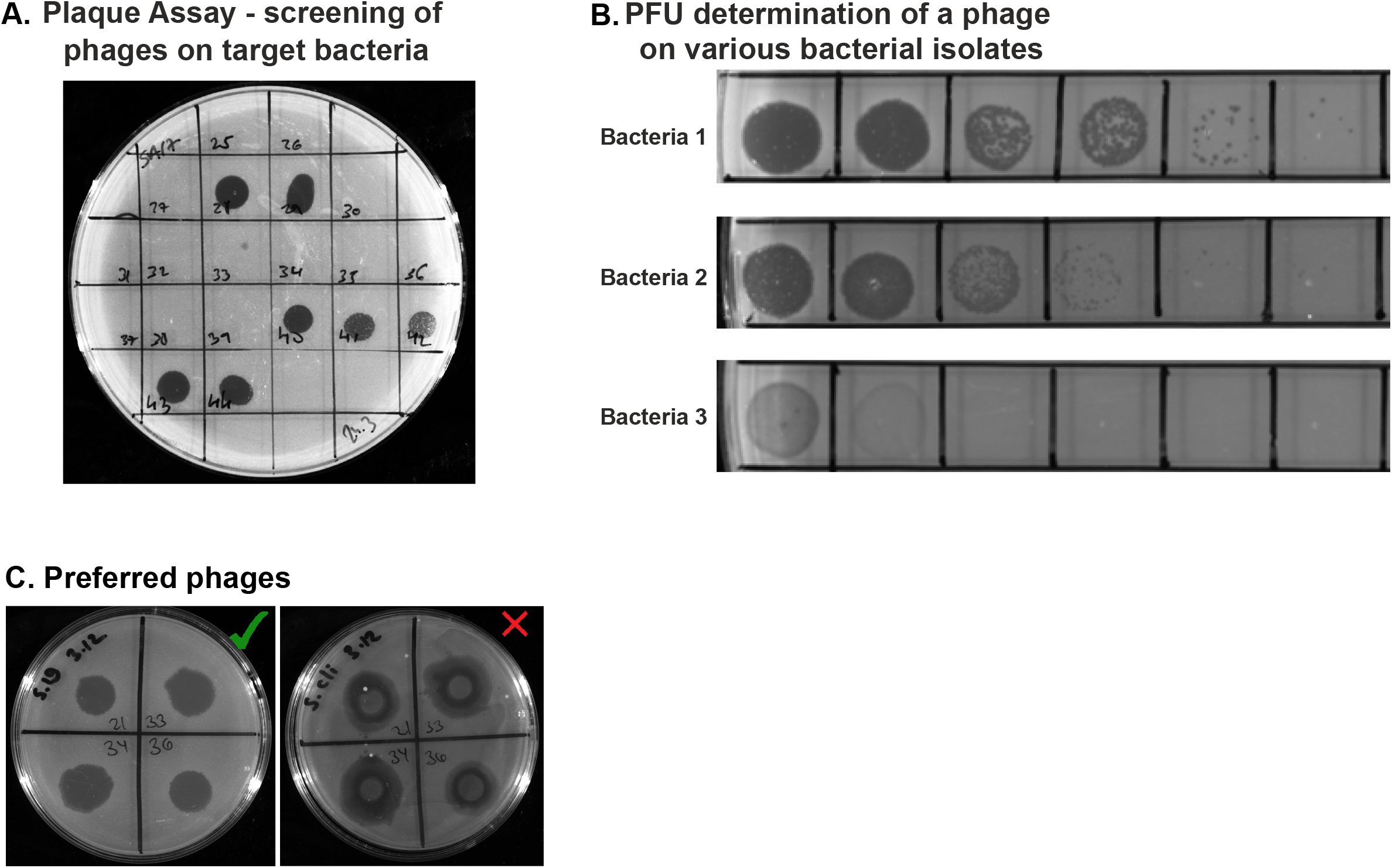
Using Plaque Assay to Match phages to the target bacteria. (A) An example of plaque assays with multiple phages, exhibiting different lytic efficacies according to plaque morphologies on the targeted bacterial isolate. For the assay, 0.2 ml of stationary bacterial culture were plated using 3 ml of agarose (0.5%) on agar plates, followed by spotting of 5-microliter drops of the tested phages and overnight incubation at 37 °C (demonstrated here by phages of *S. aureus*). (B) An example of PFU determination of the same phage on three different bacterial isolates of the same species, representing the change in phage titer and morphology according to the tested bacteria. To this end, the plaque assay was performed with phage suspensions following serial dilutions by 10-fold. The number of plaques for every given dilution can be counted, and the PFU concentration in the initial sample is accordingly calculated for each isolate (demonstrated here by a phage of *Providencia stuartii*). (C) Examples of (left) clear plaques, and of (right) turbid plaques with bacterial regrowth (demonstrated here using a phage of *Cutibacterium acnes*).

Several parameters should be considered when selecting phages for treatment using the plaque assay. As a rule of thumb, phages producing clear areas when spotted on bacterial cultures, with no bacterial colonies growing in them should be preferred (Fig 2C). Plaque size may be influenced by virion size, adsorption rate, or indicate the ability of the phage to affect non-dividing cells^27^. However, their size and morphology may also be influenced by the agar or agarose concentration in the soft layer (Fig S1), thus requiring protocol standardization among different phages^28^. Bacterial colonies within the plaques may indicate the level of pre-existing resistance against the phages^29^. Turbid plaques may indicate the presence of a lysogenic phage^21^, which may be less suitable for treatment, but not unquestionably excluded from use (see discussion).

A very important issue regarding infective phage enumeration using the plaque assay is that it depends on the tested bacteria. Phage titer and morphology may significantly vary when tested on different bacterial isolates (Fig 2B). This observation, which had been made in the early 1950s by Luria, Weigle, and Bertani^30,31^, later led Arber, Meselson, and others to discover the restriction enzymes^32^, a discovery that changed the face of molecular biology and resulted in the 1978 Nobel Prize^33^. Thus, every phage titer report should include the specific target bacterial strain on which it was tested. Moreover, PFU enumeration may be influenced by the growth stage of the bacteria, which should also be reported^34^. Final PFU results are usually determined within 24 hours, although this is dependent on the growth rate of the target host. Nevertheless, changes may occur in the plate following evaluation. New plaques can appear, and regrowth of bacteria or isolated colonies may cover the plaques. These changes should not dramatically affect the conclusion regarding the candidate phage, however, monitoring and recording of the results seen on the plates are recommended, preferably by automated methods^35^.

The titer of phages in prepared stocks may drop over time, as has been previously observed^12^. Hence, phage enumeration should be performed for the specific phage stock planned for treatment, on the most recent target bacterial isolate, as close as possible to treatment initiation.

#### B. Growth Kinetics of Liquid Cultures

Another approach used to determine phage efficacy and to test the match of phages to the bacterial target is growth kinetics monitoring (Fig 3). The main advantage of this approach is the real-time evaluation of phage-bacterial interactions. Recording the entire growth curves of bacteria in the presence of phages, rather than evaluating only the endpoint is essential, because in many cases bacterial regrowth may occur after lysis, or after prolonged growth inhibition. Furthermore, this method allows simultaneous high throughput evaluation of multiple phage-bacterial combinations and conditions such as multiple phage cocktails, phage-antibiotic combinations, and the effect of the phage(s) on polymicrobial cultures. Real-time growth kinetics monitoring may be assessed using the following techniques:

**Fig 3.**
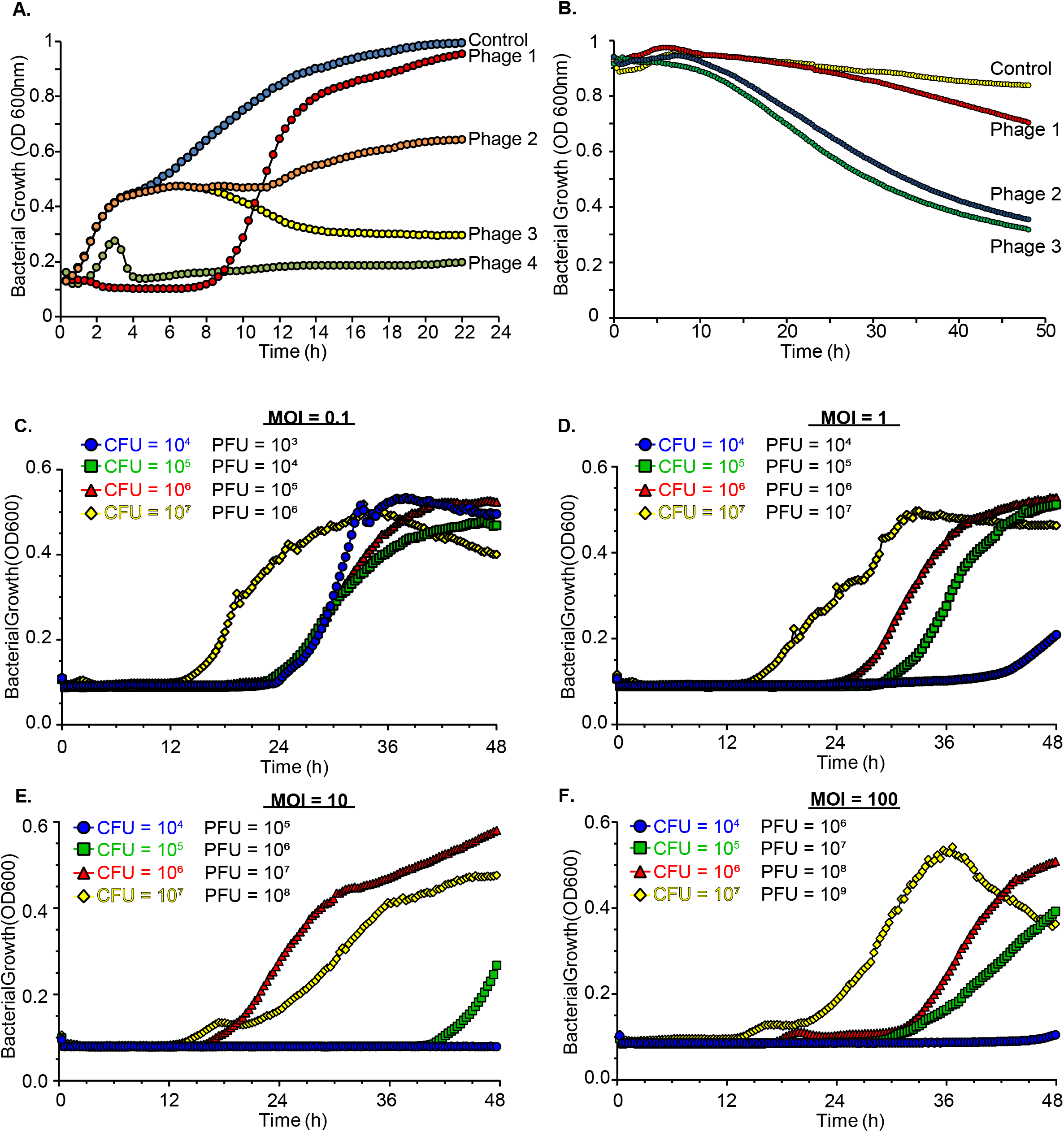
Phage Evaluation by Bacterial Growth Kinetics. Growth curves of (A) logarithmic and (B) stationary bacterial cultures, alone (control) or with the addition of different phages, determined by repeated optical density (OD_600nm_) measurements. The different phage suspensions were added to bacterial cultures in 96-well plates to a total volume of 0.2-ml per well. Optical density measurements were then recorded at 37°C with 5-sec shaking every 20 min in a 96-well plate reader at 600 nm. Potential variances in inhibitory and lytic effects of various phages are shown (demonstrated here using phages of *P. aeruginosa*). (C-F) Growth curves of bacteria treated by phages, determined by repeated OD_600nm_ measurements, with constant values of a multiplicity of infection (MOI) in every graph. Each graph illustrates different theoretical combinations of initial bacterial and phage concentrations, indicating that significant variance in bacterial growth kinetics may be seen in the same reported MOI. Of note, for lines presenting the same initial bacterial concentration (represented by the same color), similar growth kinetics are seen regardless of phage concentration or MOI (demonstrated here using a phage of *P. aeruginosa*).

##### Method 1: Cell Concentration

Measuring culture turbidity at an optical density (OD) of 600nm (Fig 3) as an indicator of cell concentration a common method for determining the growth or lysis of liquid bacterial cultures, and has been previously used in clinical phage applications^10^. The advantages of this method include its simplicity since it does not require any additional compounds except the bacteria and tested treatments. Moreover, OD measurements are dynamic and reversible, *i*.*e*., may show culture lysis even following bacterial growth (Fig 3A, yellow line), or bacterial growth following inhibition periods (Fig 3A, red line). Thus, this approach allows the assessment of phage efficacy in both logarithmic cultures in the exponential phase of growth (Fig 3A) and against non-dividing cells in the stationary phase of growth (Fig 3B). The disadvantage of this method is that it is disturbed by clumps and aggregates, causing artificial “noise” commonly seen after bacterial lysis (Fig S2) and that it does not differentiate between viable and dead cells. In addition, OD is sensitive to cell size, which may change under different conditions. Nevertheless, these changes should not limit the determination of phage efficacy.

##### Method 2: Cell Respiration

A real-time evaluation method based on a color reaction following the reduction of tetrazolium dye by cell respiration^36^ was used in several clinical phage-therapy publications ^7,8,10^. This method is less sensitive to aggregation than cell concentration. However, it depends on bacterial respiration^36^, and thus, in contrast to OD, is not suitable for the evaluation of phages against non-dividing cells, bacteria in low metabolic conditions such as stationary cultures, and bacteria under bacteriostatic inhibitory conditions. Another drawback of this method is its irreversibility due to the color reaction, which may not detect lysis following growth. Lastly, this method requires the addition of compounds and dedicated readers such as the OmniLog™instrument^36^.

##### Method 3: Culture Heat

Isothermal microcalorimetry is a real-time phage kinetics evaluation method based on the determination of heat production by bacterial cultures as an indication for growth. The main advantage of this method is its ability to assess bacterial growth and lysis in biofilm in addition to planktonic cultures. However, it requires dedicated isothermal calorimeters^37^.

#### Additional Methods for Growth Kinetics Evaluation

Various methods used to follow bacterial growth kinetics can be adopted for phage efficacy determination. Examples include methods based on flow cytometry, on methylene blue dye reduction, or the length of bacterial lag phase demonstrated by optical density^38–40^.

##### Logarithmic or Stationary Cultures?

Phages and antibiotics may present different efficacies against dividing logarithmic bacteria in comparison to stationary bacteria. It is difficult to determine which of these growth stages *in-vitro* better represents the clinical settings, influenced by variables such as the site and the acuity of infection. Thus, we recommend testing simultaneously the efficacies of candidate phages on both overnight bacterial cultures (stationary, Fig 3B) and diluted cultures (logarithmic, Fig 3A). Phages that show high efficacy in both stages should be preferred.

##### Multiplicity of Infection (MOI)

An important open question remains regarding the concentration of bacteria and phages used *in-vitro* that most accurately predict treatment outcomes. Usually, when bacteria and phages are mixed, the multiplicity of infection (MOI), *i*.*e*. the ratio of phages to bacteria, is described. However, data further suggest that a description of MOI alone is not sufficient, as the absolute concentration, particularly of bacteria, can significantly affect the results^41^. Thus, the efficacy of a given phage on a specific bacterial strain may dramatically vary if the bacterial concentration is changed, even under identical PFU or MOI in every sample (Figs 3C-F). To overcome this problem, we believe it is important to test the bacteria in several starting concentrations.

##### Viable Cell Count Following Kinetic Assays

The last step in growth kinetic evaluation should be a bacterial-colony-forming unit (CFU) enumeration, as an estimate of the lethality of the tested phage. The growth curve can occasionally be inaccurate, especially when bacterial concentration is very low, around the detection limit. Thus, the viable bacterial count may be used as a validation tool, allowing phages exhibiting increased lethality to be chosen. However, it is important to mention that when samples are plated for bacterial enumeration, phages may be carried on and interfere with the colonies grown on the plate, therefore disrupting the results. Hence, it is recommended to wash the cells before plating.

##### Scoring Phage Efficacy

Phage evaluation based on growth kinetics should be automated and standardized for the quantitative assessment using reproducible scoring algorithms. These algorithms should take into account several parameters including length of lag phase, maximum growth, the area under the curve, the slope of lysis, and regrowth following lysis. Additionally, they may take into consideration the plaque assay, with parameters such as plaque titer, morphology, and size. Several scoring methods for phage evaluation have been previously suggested, such as The Virulence Index^42^, assessing the virulence of phage by comparing the integrated area of bacterial reduction curves to the integrated area of the phage-free controls. Constant improvement and implementation of similar algorithms will allow the standardization of quantitative phage evaluation.

#### C. Phage-Antibiotic Combinations

The interactions between bacteria, phages, and antibiotics can be unpredictable and vary between synergistic, additive, neutral and antagonist effects^43–46^, studied from as early as 1945^47^. Examples of phage-antibiotic interactions are presented in Fig 4A. The positive effects between these agents can be explained by several mechanisms, ranging from merely reduced bacterial density posed by the treatment combination, to direct negative interactions between resistance mechanisms of different agents^48^. Moreover, a method of deliberate use of phage resistance has been previously proposed, termed Phage Steering, relying on trade-offs between bacterial resistance to phages and antibiotics^49^.

**Fig 4.**
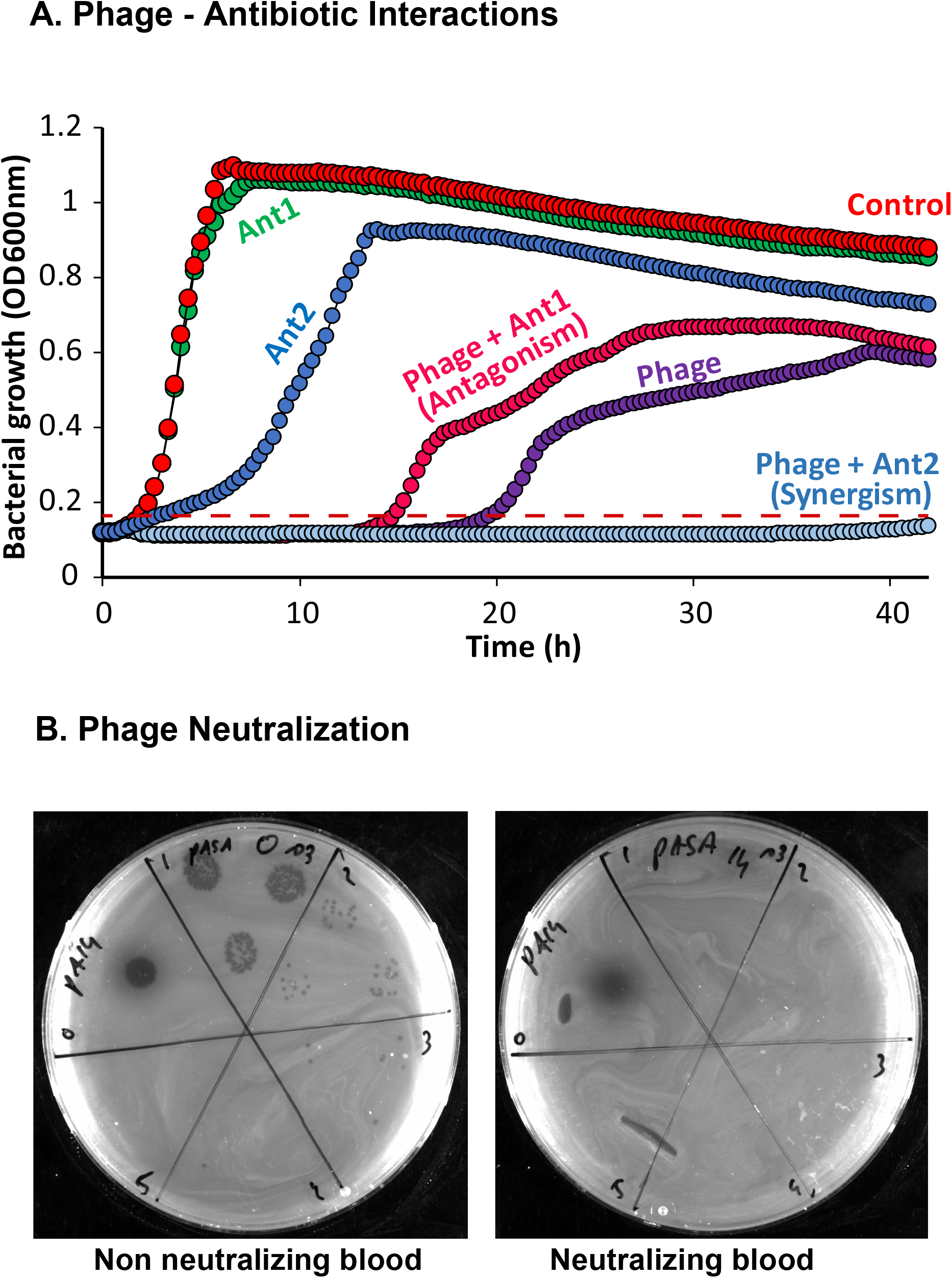
Phage-Antibiotic Checkerboard and Phage Neutralization Assays.. (A) Phage-Antibiotic Checkerboard. Growth curves of bacteria either alone, or treated with phages, antibiotics, or their combinations. The tested treatments were added to bacterial cultures in 96-well plates to a total volume of 0.2-ml per well. Optical density measurements were then recorded at 37°C with 5-sec shaking every 20 min in a 96-well plate reader at 600 nm. Different effects of phage-antibiotic combinations can be seen, including synergism (light blue line) or antagonism (red line), demonstrated by prolongation or shortening of the lag phase, respectively (demonstrated here using a phage of *E. faecalis*). (B) Phage Neutralization. Diluted phage suspensions were added to different blood samples drawn from a patient treated by the phage. One sample (left panel) was drawn before treatment initiation, and the second (right panel) was drawn following two weeks of treatment. The mixed samples were incubated for 15 min at 37°C in a drum rotator at 10RPM, followed by 30 min incubation at room temperature without shaking. The samples were then serially-diluted and spotted on bacterial cultures as described for PFU determination. It can be seen that in the right panel the PFU significantly decreased compared with the left panel, indicating phage neutralization by this sample (demonstrated here using a phage of *P. aeruginosa*).

Many of the patients recently treated by phage therapy have received concomitant antibiotic regimens^50^. However, the current clinical knowledge regarding such combinations is still not sufficient and is based mostly on sporadic observations^7,10,45^.

Based on the above, in our view, it is highly important to specifically test the efficacy of the planned combination for every phage treatment before initiation. The antibiotics selected for testing should initially be based on standard sensitivity results^3^ and clinical judgment. These tests can be performed using Phage-Antibiotic Checkerboard (PACB), allowing simultaneous analysis of various phage-antibiotic combinations, performed in 96-well-plates, based on bacterial growth kinetics monitoring (Fig 4A, Fig S3). Such testing can be completed within 24 hours for fast-growing pathogens, thus allowing to choose the most efficient treatment combination without a significant prolongation of the “time to treatment”.

An example of such a checkerboard can be found in our previous work ^10^, where we have simultaneously assessed the combined effect of two lytic phages and several different antibiotics on *Acinetobacter baumannii* and *Klebsiella pneumoniae*, isolated from polymicrobial osteomyelitis. This analysis helped us identify the phage-antibiotic combination associated with the best inhibitory effect, which was later used for successful treatment^10^.

#### D. Phage Neutralization by Immune System

Anti-phage activity mediated by the humoral response has been repeatedly observed during or after phage therapy^7,51,52^. Furthermore, it was shown that antibodies against the phage T4 exist in the majority of the healthy population^53^. We speculate that the chance to find anti-phage antibodies in patient sera even increases in cases of prolonged chronic infections. Nevertheless, anti-phage antibodies do not necessarily exclude sufficient treatment outcomes^54^, and the level of the humoral response may vary depending on the phage-type^52^. Thus, at this time it is unclear if the presence of antibodies against phages should lead to exclusion of the phage from treatment. Yet, in our view, it is useful to test patients for phage neutralization, by mixing the phage solution with the patient’s serum and performing PFU enumerations following incubation^54^, as presented in Fig 4B. In the case of positive phage neutralization, the tested phage should have lower priority in treatment design.

##### Our suggested recommendations

1. ***Assessment of bacterial susceptibility to phages should be performed before any treatment***.
2. ***Evaluation of phage efficacy should be carried out using at least two simultaneous methods, including the plaque assay and a liquid growth kinetics monitoring technique***.
  2.1. ***Plaque assay: phages producing clear plaques in high PFU on the target bacteria should be preferred***.
  2.2. ***Liquid growth kinetics: the assay should be performed on both logarithmic and stationary cultures followed by viable cell count validation***.
3. ***The combined efficacy of the phages intended for the treatment and the clinically preferred antibiotics should be tested on the target bacteria by bacterial growth kinetics monitoring (Phage-Antibiotic Checkerboard)***.
4. ***Patient sera should be assessed for the presence of specific anti-phage antibodies, and if possible, phages not undergoing neutralization should be preferred***.
5. ***Phages considered for treatment should be routinely checked for PFU on the targeted bacterial isolate***.

#### Deep Screen and Special Cases

Thus far we have discussed our suggested pipeline for phage matching to bacterial targets, performed before phage therapy to allow the most suitable treatment design. However, occasionally, clinical cases may require additional laboratory considerations and a deeper screen, aiming to enhance the rate of treatment success. In this chapter, we discuss our approach and suggestions for troubleshooting pitfalls or special cases.

##### A. Urgency in Phage Application

For most fast-growing bacteria, which comprise the vast majority of current phage applications, the results for phage matching to bacterial targets using both the plaque assay and the liquid growth evaluation methods are usually examined within 18 hours (overnight). Nevertheless, in clinical cases of an emergency, results regarding phage efficacy from both methods may be obtained within only a few hours, thus significantly accelerating the next steps towards phage application (Fig 5). In this case, it is even more important to establish the results using several simultaneous methods to increase confidence and to later verify the results with overnight cultures. Thus, an initial accurate, albeit suboptimal, treatment can be provided within a few hours for fast-growing bacteria.

**Fig 5.**
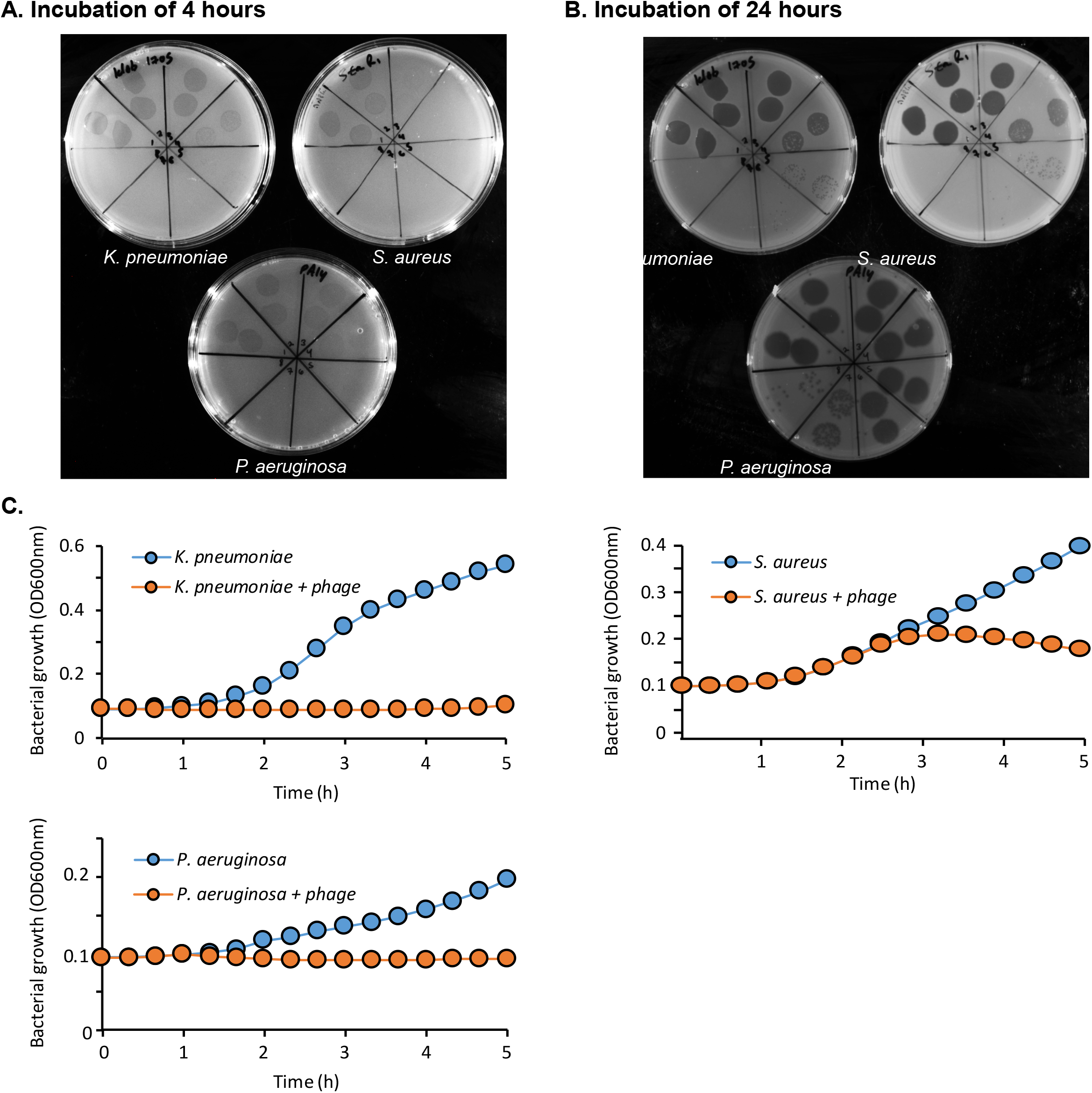
Rapid Phage Screening. (A-B) Plaque Forming Unit (PFU) enumeration of three phages on different bacterial strains, determined by spotting phage suspensions following serial dilutions by 10-fold. The plates were incubated at 37°C for (A) 4 hours and (B) 24 hours, demonstrating the ability to predict phage efficacy following short incubation periods. (C) Growth curves of different bacterial strains, alone or treated by phages, determined by optical density measurements recorded at 37°C with 5-sec shaking every 20 min in a 96-well plate reader at 600 nm. Phage effect on the bacterial growth kinetics can be observed within only a few hours of incubation (demonstrated here using phages of *K. pneumoniae, P. aeruginosa*, and *S. aureus*).

##### B. Polymicrobial Infections

Polymicrobial infections represent special, hard to treat clinical conditions^55^. Certain bacterial strains may provide resistance abilities to the other strains^56^, emphasizing the importance of their concomitant eradication^10^. Before phage treatment of polymicrobial infections, it is recommended that all intended treatments be tested together on the mixed bacterial cultures by bacterial growth kinetics monitoring, for prediction of the most suitable treatment regimen.

##### C. Biofilm

Biofilms are another significant clinical challenge, characterized by persistent infection that is difficult to treat with current antimicrobial agents^57,58^, which may be overcome by therapy with phages that have biofilm activity^37,45,59^. This is, perhaps, due to the proximity of the cells in the biofilm, which enhances phage infection. However, combined therapies of several phages or additional antimicrobials may be needed for effective biofilm control^60^, and thus should be properly selected. This was demonstrated by Chan *et. al*. in the case of *P. aeruginosa* biofilms, treated with either phage, antibiotics, or their combination before clinical application^45^. Numerous methods have been described for the evaluation of biofilm eradication by phages, including crystal violet staining^59,61^, confocal microscopy^59^, and electron microscopy^44^. However, many of these methods require relatively long periods necessary for biofilm formation, lasting at least a couple of days. Therefore, we recommend testing for phage efficacy using such models only in clinical cases involving biofilm, which also allows a delay in time to treatment, such as prolonged chronic infections.

##### D. An Individual Phage or a Cocktail?

Phage therapy is commonly applied in the form of “phage cocktails”, *i*.*e*., concomitant administration of several phages targeting either individual or various bacterial strains ^11,62^. The logic behind the use of phage cocktails arises from the idea that simultaneous treatment targeting a variety of bacterial receptors with diverse antibacterial pathways will more efficiently decrease the bacterial burden and the development of resistance^63^. Simultaneous administration of phages was found in several *in vitro* and *in vivo* models superior or equal to their sequential application^64^. Nevertheless, this question requires more hypothesis-driven studies.

Phage cocktails can be either “fixed”, consisting of pre-designed phage combinations^13^, or custom made per case^62^. From a regulatory point of view, “fixed “cocktails with relatively wide coverage, marketed as shelf products, may be considered safer than *ad-hoc* preparations^65^. Nevertheless, the complex relationship between bacteria and multiple phages may negatively impact treatment efficacy^66^, for example, due to possible antagonism or development of multi-resistance.

Accordingly, in our approach, the use of phage cocktails may be especially required if no phage has shown sufficient efficacy when tested individually^67^, or in cases of polymicrobial infections^10^. Note that before cocktail preparation, all intended treatments should be tested together using a Phage Checkerboard based on bacterial growth kinetics monitoring. Moreover, to avoid changes in the composition of the cocktail due to differences in phage properties, we suggest combining the phages as close as possible to their administration.

##### E. Method Discrepancy

In the case of discrepancy between different evaluation techniques ^68^, such as the plaque assay and the liquid growth kinetics, or between logarithmic and stationary growth stages, phage cocktails with increased coverage in all methods ^67^ should be considered.

##### F. Antibiotic Mismatch

In cases when all the tested phage-antibiotic combinations are not efficient against the target bacteria, *e*.*g*. due to interference between these two agents, non-intuitive antibiotics should be also screened. For instance, even antibiotics against which the target bacterial strain is resistant *in-vitro* may synergize with a phage. Such a synergistic effect has been recently shown *in–vitro* ^44^ and in clinical settings ^7^.

###### Our suggested recommendations

1. ***In cases of emergency, results regarding phage efficacy may be obtained within a few hours and later verified following overnight growth***.
2. ***In cases of polymicrobial infections, optimal treatment should be initially individually matched to each bacterium, followed by an evaluation of the combined mixture of all bacterial strains and intended treatments (all phages and antibiotics)***.
3. ***In cases involving biofilm, the efficacy of the intended treatment combination should be tested using the biofilm model***.
4. ***Single phages should be preferred over phage cocktails unless individual phages have not shown sufficient efficacy in all phage matching methods***.
  4.1 ***All phages intended to be combined with a phage cocktail should be tested together on the target bacteria by bacterial growth kinetics monitoring (Phage Checkerboard)***.
  4.2 ***When using a phage cocktail, the different phages should be combined into a cocktail as close as possible to the treatment initiation***.
5. ***In cases of phage-antibiotic mismatch, screening of antibiotics for potential combination with phages may be expanded beyond the standard bacterial susceptibility results***.

#### Monitoring during Treatment

The role of Clinical Phage Microbiology may not end with the initiation of the treatment, and monitoring should continue throughout the treatment. The main point to be addressed in this stage is the potential emergence of bacterial resistance to the applied treatments, as previously seen^7,8^. For example, Schooley *et al*.^7^ have described the isolation of a bacterial strain resistant to two initially used phage cocktails (each comprised of 4 phages), within 8 days after their introduction, leading to the application of additional phages. In such cases, new matching steps including phage screening should be performed and the treatment should be changed accordingly, because of changes in bacterial properties ^7^.

##### Resistance Simulation and Preparedness

Since the number of mutations leading to bacterial resistance is not infinite, and some are more likely to develop, we can deliberately induce *in-vitro* resistance, to simulate their appearance and match suitable treatments to these isolates. Accordingly, in case that resistance emerges during treatment, there is a high probability that suitable treatments would be immediately available. Resistant mutants can be induced by growing the bacterial strain in the presence of the used phage for several rounds or by testing colonies growing on phage plaques^29^. We had previously demonstrated this technique by *in-vitro* inducing resistance for a bacterial isolate from a patient treated by phage therapy and isolated a new phage efficient against it^10^. Fortunately, in this case, no resistance developed in the patient.

###### Our suggested recommendations

1. ***In cases of bacterial isolation from a patient during treatment, the new isolates should be tested for susceptibility to the administered treatment. Accordingly, new steps for phage matching should be performed for treatment adjustment***.
2. ***Phage resistant bacterial mutants may be deliberately induced in-vitro, for prompt treatment adaptation in case of resistance***.

## Discussion and Future Directions

Here we present a new concept, Clinical Phage Microbiology, aimed to establish laboratory guidelines supporting the clinical use of phage therapy. This concept takes into account the variability among phages, stressing the need for individual laboratory evaluation of the specific suitability of every phage to the targeted bacterial strain and the clinical settings. Assumptions regarding phage efficacy should not be made solely according to general concepts without supporting laboratory evidence.

We have emphasized that phage selection should be performed using a combination of several recognized techniques, with an agreement in their results, and not according to theoretical compatibility. Every relevant phage from the local phage banks containing characterized and purified phages should be evaluated. Furthermore, following genome sequencing, it may be possible to identify and exclude phages containing toxin-producing genes, virulence factors, or antimicrobial resistance genes ^15^. Phage-associated toxins and virulence factors have been linked to various infectious diseases ^69,70^, highlighting the need for proper genetic evaluation of phages intended for treatment. However, as opposed to the generally-reported approach, phages containing genes associated with a lysogenic phase should not be unavoidably excluded

^71^. This is because the phage life-cycle, *i*.*e*. lysogenicity, may be isolate-specific, and should be experimentally defined for the specific phage-bacterial interaction due to potential lytic variants. This is particularly important in cases where no phages with an obligatory lytic cycle can be found, thus allowing to expand of the search to phages that potentially represent natural lytic variants, as previously achieved via phage-engineering ^21^.

Following phage selection, suitable treatment protocols can be constructed by the suggested practices. This includes the evaluation of the combined effect of the phages with additional antimicrobials, the specific infection settings, and the patient’s immune response to the treatment. Lastly, laboratory methods for follow-up during therapy and preemptive preparedness for resistance are suggested, enabling real-time clinical adjustments.

Several points remain currently unaddressed, requiring further investigations. How should phage dosage and timing be determined? Should phages be administered simultaneously to antibiotics, or are there beneficial effects to specific treatment order selection? Are there changes in phage pharmacokinetics during treatment progress? How can potential interactions between phages and other commonly used medications be assessed? Only sufficient data collected from current cases and animal studies will allow better address these and additional issues, essential for treatment design. The last and perhaps the most important step of Clinical Phage Microbiology is a thorough data collection from every phage application. A methodological structured collection of data regarding the bacterial isolates, the available phenotypic and genotypic data of the phages, the influence of all tested antibiotics on phage efficacy, and all relevant clinical information concerning the patients and treatment outcomes should be recorded and appropriately labeled. This will later allow the creation of comprehensive datasets that can be used to detect the effects of different features on phage selection and treatment design. Accordingly, artificial intelligence (AI) will lead to an even more accurate selection of phages for every patient, based both on genetic predictors and additional detected relationships. Furthermore, novel technological developments, such as accessible techniques for phage purification will increase the potential of phage engineering or training, for the most accurate adaptation of phages for every patient.

In summary, it has taken clinical microbiologists five decades to work out the current framework for antibiotic selection using *in vitro* assays, in an iterative process, in which laboratory innovations were tested in the clinic, and clinical success and failure were correlated with laboratory results. The same approach will need to be taken as we develop the principles of Clinical Phage Microbiology. The addition of Clinical Phage Microbiology, as suggested here, to the standard procedures performed by departments of clinical microbiology is a vital step towards the integration of phage therapy into the clinical practice.

## Acknowledgments

The authors would like to thank the “United States - Israel Binational Science Foundation (BSF)” grant #2017123, the Israel Science Foundation (ISF) IPMP grant #ISF_ 1349/20, the Rosetrees Trust grant A2232, and the Milgrom Family Support Program.

## Figure legends

**Fig S1. The Effect of Agarose Concentrations on Plaque Appearance**. An example of different plaque morphologies according to agarose concentration. A plaque assay was performed on agar plates with different agarose concentrations (left plate - 0.5%, right plate - 0.3%, and bottom plate – 0.15%), by spotting three tested phages (A, B, and C) followed by overnight incubation at 37 °C. It can be seen that the agarose concentration has a noticeable effect on the clarity of the lysis zones (demonstrated here by *Serratia marcescens)*.

**Fig S2. Optical Density Noise**. Bacterial growth curves are determined by repeated optical density (OD_600nm_) measurements. It can be seen that the curves show artifacts originating from probable aggregates or clumps, especially during bacterial lysis, disturbing graph interpretation.

**Fig S3. Optical Density Checkerboard**. Illustration of checkerboard performed by a 96-well plate optical reader, presenting the ability to simultaneously assess multiple different phage-antibiotic combinations by optical density (OD_600nm_) measurements. Each assay is performed in triplicate, shown by the blue lines, and compared to controls, shown in red. The outermost layer is intentionally filled with distilled water to avoid fluid dehydration at 37°C incubation.

