## Supplementary figures and images for "Clinical Phage Microbiology: A suggested *in-vitro* framework for phage therapy"

### Fig S1

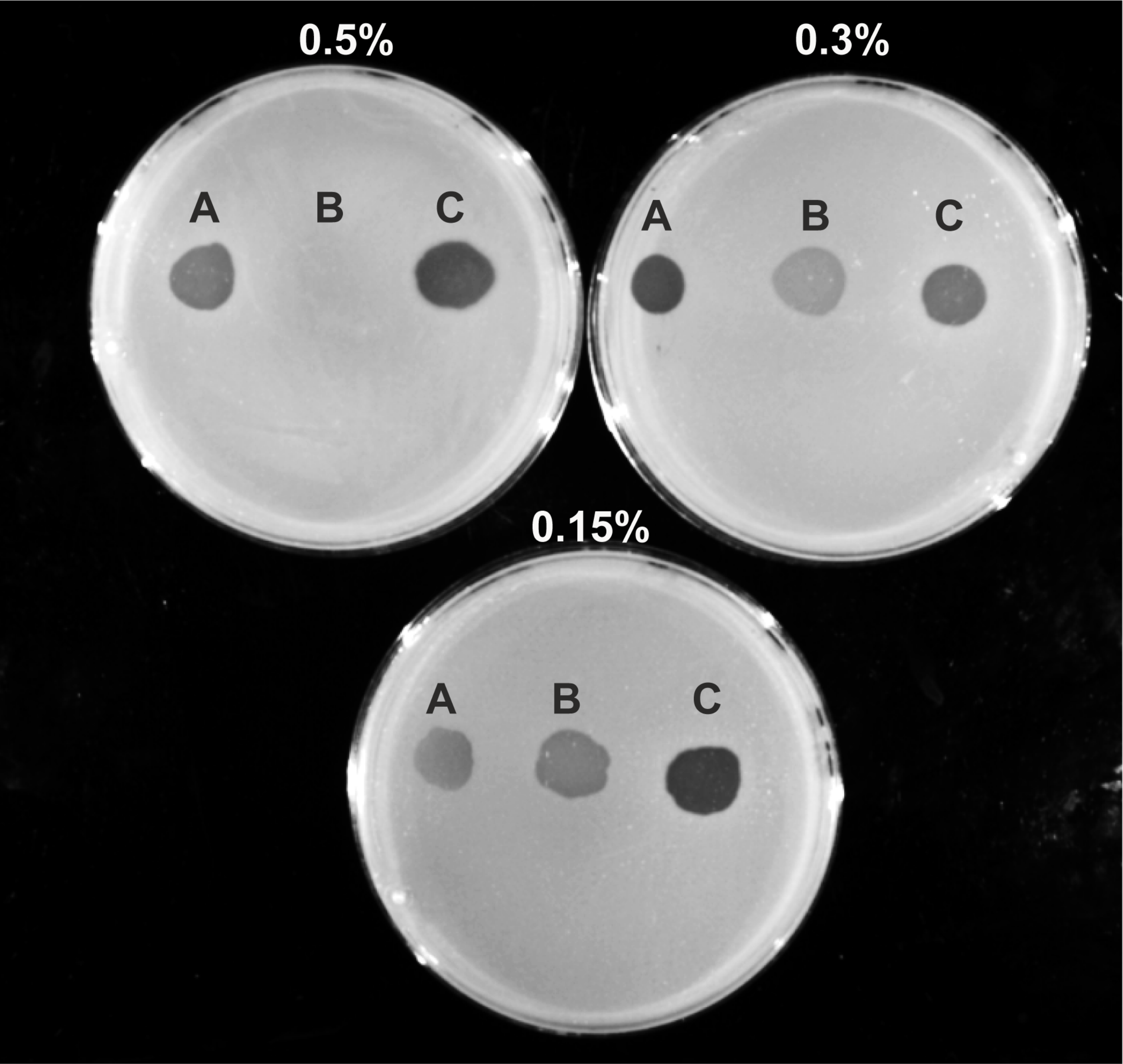

### Fig S2

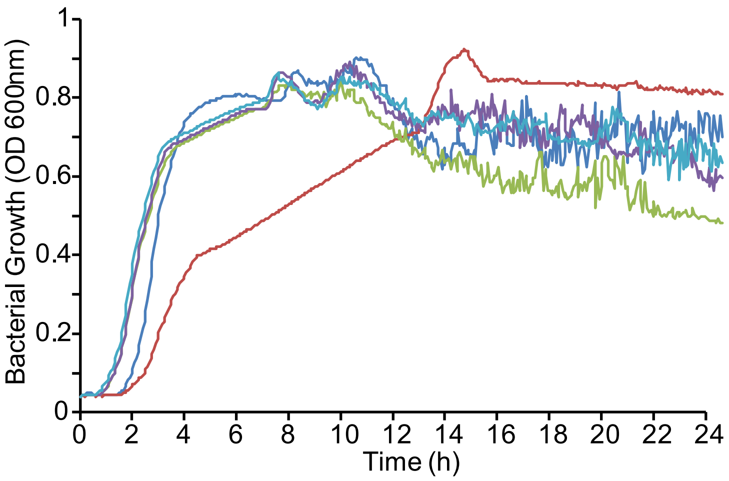
