## Supplementary material for "Clinical Phage Microbiology: A suggested *in-vitro* framework for phage therapy": FigS3

**Checkerboard**

| | 1 | 2 | 3 | 4 | 5 | 6 | 7 | 8 | 9 | 10 | 11 | 12 |
|---|---|---|---|---|---|---|---|---|---|---|---|---|
| A | | | | | | | | | | | | |
| B | | | | | | | | | | | | |
| C | | | | | | | | | | | | |
| D | | | | | | | | | | | | |
| E | | | | | | | | | | | | |
| F | | | | | | | | | | | | |
| G | | | | | | | | | | | | |
| H | | | | | | | | | | | | |
